# Ferroptosis is determined by chloride ions

**DOI:** 10.1101/2023.02.07.526847

**Authors:** Dichun Huang, Zhangshuai Dai, Chao Wang, Haiying Mai, Junyi Zhao, Lingbo Cao, Sijie Wang, Zhun Dai, Kun Xu, Ronghan He, Shaoshen Zhu, Yanchen Chen, Tiejun Bing, Xinzhou Zhu, Ting Gang Chew, Junqi Huang

**Author notes:** These authors contributed equally to this work (Dichun Huang, Zhangshuai Dai, Chao Wang). These authors jointly supervised this work (Xinzhou Zhu, Ting Gang Chew, Junqi Huang).

## Abstract

Ferroptosis has emerged as a deliberate type of programmed cell death that manifests marked importance ubiquitously in health and diseases. However, after a decade of research, the mechanisms of ferroptosis execution remain unclear. Here we identify chloride ions (Cl^-^) as essential determinants of ferroptosis. Water homeostasis manipulated by extracellular solute concentration disrupts ferroptotic cell death. Hyperosmotic stress attenuates ferroptosis and endues cells with high lipid peroxidation. Analyses of a fluorescent chloride probe show that Cl^-^ fluxes into the cytoplasm during ferroptosis, substantiating a role for Cl^-^ to drive water flow. Depletion of extracellular chloride ions ([Cl^-^]_o_) from culture media congruously confers resistance to ferroptosis. The [Cl^-^]_o_-depleted ferroptotic cells fall into two populations: cells with low lipid peroxidation; cells with high lipid peroxidation but not cell swelling or cell rupture. Contrarily, solitary [Cl^-^]_o_ overload is sufficient to elicit ferroptosis without canonical ferroptosis inducers. Further experiments show that ferroptotic cells depolarize and [Cl^-^]_o_ is positively correlated with this process. Membrane depolarization upregulates the level of lipid peroxidation, suggesting that membrane potential may be a universal mechanism governing ferroptosis. Together, our findings reveal that ferroptosis is determined by chloride ions.

## Introduction

Coined in 2012, ferroptosis is well-known as a type of iron- and lipid peroxidation-mediated programmed cell death^1,2^. Since its inception, emerging evidence has convincingly demonstrated enormous potential of ferroptosis in treating chemotherapy resistant cancer types and organ damages^3–15^. Understanding the molecular mechanisms governing ferroptosis is therefore a prerequisite for better clinical application.

Morphologically, cell swelling and eventual ruptures of the plasma membrane are prominent features of ferroptosis, irrespective of the inducers^16,17^. At the molecular level, irons, proteins and metabolites such as system x_c_^-^, GPX4, p53, FSP1, YAP, oxidoreductase POR, dihydroorotate dehydrogenase (DHODH), cyst(e)ine, glutathione (GSH), reactive oxygen species (ROS), polyunsaturated fatty acids (PUFAs), vitamin K and lipid peroxides (oxidative modification of phospholipids) are prevailing players in ferroptosis^9,18–27^.

Ferroptosis has so far been most studied in pathways controlling the generation of lipid peroxidation. The execution mechanism of ferroptosis has long been disputed. Several studies implied a plausible sandwich pathway between lipid peroxidation and the execution of ferroptosis cell death^16,28–30^. However, this plausible ferroptotic downstream mechanism remains unclear and an anticipatory cell stage at which ferroptotic cells survive with/through high lipid peroxidation over an extended period was not disseminated. Currently, it is widely considered that unrestrained lipid peroxidation directly ruptures the plasma membrane in this type of unique cell death modality, presumably without the participation of downstream proteins. This is so far evaluated as a sharp difference between ferroptosis and other programmed necrotic cell death, some of which are mediated by the pore-forming assemblies made up of MLKL or the gasdermin family proteins^31–37^. These pore-forming proteins perforate the plasma membrane, which promotes water inflow and subsequent cell rupture. However, evidences substantiating that peroxidation-distorted lipid membrane directly detonates ferroptosis in live cells is indeed long missing. Thus, the precise genuine mechanism underlying how ferroptotic cells die has largely remained unknown.

## Results

### Extra sorbitol/sucrose attenuates ferroptosis and endues cells with high lipid peroxidation

HT-1080 human fibrosarcoma cells swell prior to cell lysis in the presence of RSL3 (Fig.1a), suggesting that water influx happens and results in the plasma membrane rupture during ferroptosis. To test if this was the case, we treated cells with an increasing concentration of sorbitol (So) or sucrose (Su), which does not cross plasma membrane easily and thus lessens water from flowing into cells. Upon RSL3 treatment in the presence of sorbitol or sucrose at high concentrations, we observed that a significantly less number of cells underwent ferroptotic cell death compared to cells grown without these solutes (Fig.1b-d). The suppression of cell death by these solutes was also observed in ferroptosis induced by erastin or cumene hydroperoxide (Fig.1e-f).

**Fig. 1.**
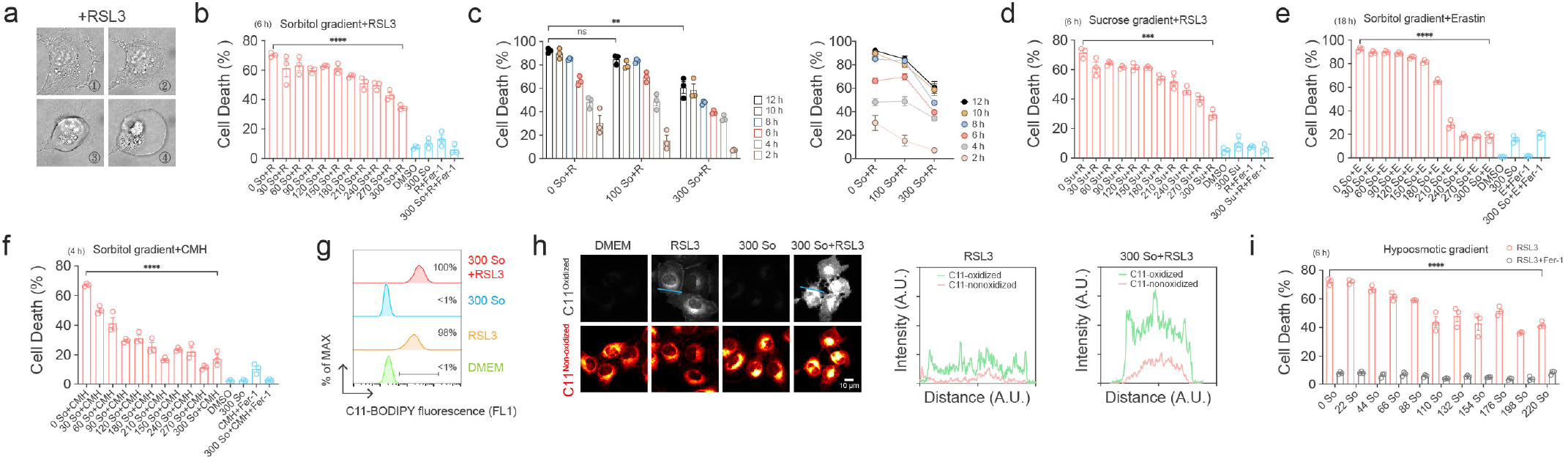
Extra sorbitol/sucrose attenuates ferroptosis and endues cells with high lipid peroxidation. **a,** Time-lapse DIC images of a RSL3-induced ferroptotic cell. Time sequence, ①<②<③<④. **b,** High sorbitol attenuates RSL3-induced ferroptosis. Sorbitol concentration, mM. **c,** High sorbitol attenuates RSL3-induced ferroptosis over time. Sorbitol concentration, mM. **d,** High sucrose attenuates RSL3-induced ferroptosis. Sucrose concentration, mM. **e,** High sorbitol attenuates erastin-induced ferroptosis. Sorbitol concentration, mM. **f,** High sorbitol attenuates CMH (cumene hydroperoxide)-induced ferroptosis. Sorbitol concentration, mM. **g,** Lipid ROS production of 300 mM Sorbitol+RSL3 cells assessed by flow cytometry using C11-BODIPY. So, sorbitol. 300 So, 300 So+RSL3 treatments were assayed at 1 h. RSL3 treatment was assayed at 2 h. **h,** Lipid ROS production of 300 mM Sorbitol+RSL3 cells assessed by confocal microscopy using C11-BODIPY. So, sorbitol. **i,** Hypotonic media created by depletion of NaCl (~110 mM in DMEM) and subsequent supplementation with gradients of sorbitol. So, sorbitol. Sorbitol concentration, mM. 0 So illustrates the NaCl-depleted media which is osmotically imbalanced. 220 So illustrates the NaCl-depleted 220 mM sorbitol-supplemented media which is osmotically balanced.

Interestingly, despite attenuating ferroptosis, these cells accumulate a high level of lipid peroxidation. Both flow cytometry and microscopy analyses showed that cells accumulated high level of oxidized C11-BODIPY when treated with RSL3 and high concentration of sorbitol, which is comparable to cells treated with RSL3 alone (Fig.1g-h). Conversely, when cells were exposed to a lower solute media (osmotically imbalanced) by depletion of extracellular NaCl, more cells underwent ferroptosis compared to cells grown in osmotically balanced media (Fig.1i). Taken together, our data suggest that ferroptotic cell death can be antagonized by elevating extracellular solutes, which potentially lowers water influx yet endues ferroptotic cells with high lipid peroxidation.

### Depletion of [Cl^-^]_o_ from culture media impedes ferroptosis

We speculated that the influx of extracellular ions towards the concentration equilibrium can contribute to water uptake thereby regulating ferroptosis. When cells are grown in osmolarity-balanced ion-free water, cell death induced by RSL3 is attenuated as compared to that of when cells grown in the PBS saline buffer (phosphate buffered saline, 154.00 mM NaCl, 5.60 mM Na_2_HPO_4_, and 1.06 mM KH_2_PO_4_) containing mostly NaCl (Fig.2a). Since other saline buffers containing NaCl also support ferroptosis execution (Fig.2b), we asked whether NaCl functions in ferroptosis. Interestingly, ferroptotic cell death is positively correlated with NaCl concentration (Fig.2c). NaCl appeared to exert its effects independent of osmotic pressure as addition of 150 mM NaCl compared to 300 mM sorbitol did not suppress ferroptosis (Fig.2d).

**Fig. 2.**
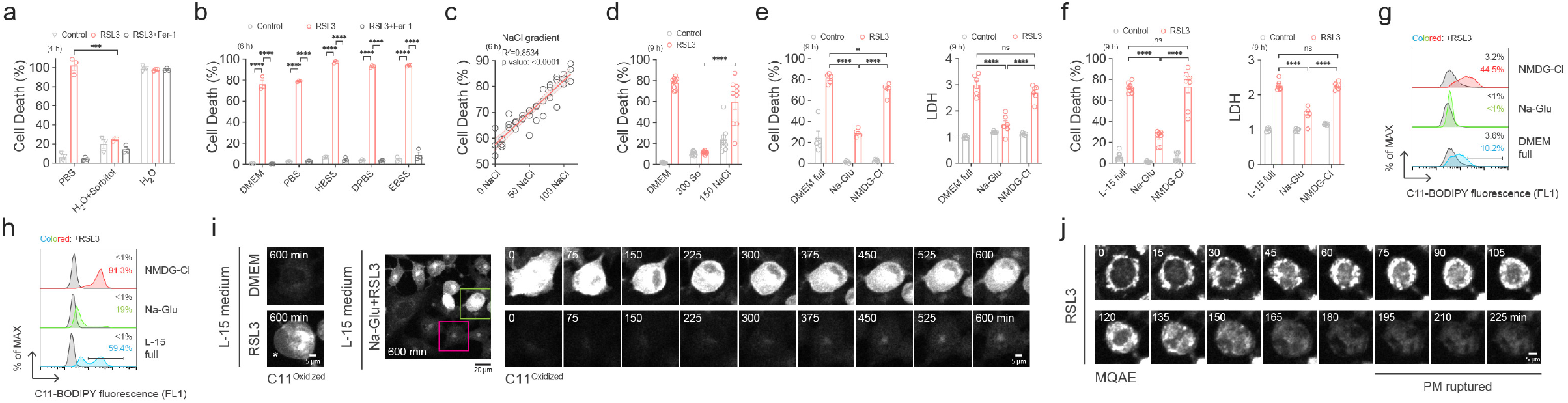
Low [Cl^-^]_o_ is capable of ferroptosis blockage. **a,** Deprivation of all extracellular ions attenuates ferroptosis. Osmolarity is balanced by supplementation of sorbitol. **b,** Ferroptosis induction in different saline buffers. **c,** Ferroptotic cell death is positively correlated with NaCl concentration in DMEM. NaCl in DMEM is replaced reciprocally by gradients of sorbitol. **d,** Isotonic DMEM media containing extra 300 mM sorbitol or 150 mM NaCl exhibit different effects on ferroptosis. **e,** Replacement of [Cl^-^]_o_ in DMEM with sodium gluconate, but not NMDG-Cl, suppresses ferroptosis. Cell death was measured by PI/Hoechst staining or LDH release. DMEM full represent complete DMEM media. **f,** Replacement of [Cl^-^]_o_ in L-15 media with sodium gluconate, but not NMDG-Cl, suppresses ferroptosis. Compared to NaCl-depleted DMEM, NaCl-depleted L-15 media contains less sodium and chloride ions. Cell death was measured by PI/Hoechst staining or LDH release. L-15 full represent complete L-15 media. **g**, Lipid ROS production in NaCl-replaced DMEM media assessed by flow cytometry using C11-BODIPY. All treatments were assayed at 3 h. Na-Glu represents NaCl-depleted Na-Glu supplemented osmolarity-balanced DMEM media. NMDG-Cl represents NaCl-depleted NMDG-Cl supplemented osmolarity-balanced DMEM media. **h,** Lipid ROS production in NaCl-replaced L-15 media assessed by flow cytometry using C11-BODIPY. 0.2 μM RSL3 was used in this experiment. All treatments were assayed at 3 h. Na-Glu represents NaCl-depleted Na-Glu supplemented osmolarity-balanced L-15 media. NMDG-Cl represents NaCl-depleted NMDG-Cl supplemented osmolarity-balanced L-15 media. **i,** Lipid ROS production in NaCl-replaced L-15 media assessed by time-lapse confocal microscopy using C11-BODIPY. RSL3 treatment alone showed cell swelling at 600 min (white asterisk). The upper right montage shows a representative NaCl-depleted Na-Glu supplemented cell with high lipid ROS throughout the course of imaging but not cell swelling or cell rupture (same cell in green square). The lower right montage shows a representative NaCl-depleted Na-Glu supplemented cell with low lipid ROS (same cell in purple square). **j,** Time-lapse confocal microscopy images of MQAE staining in a RSL3-induced ferroptotic cell. PM, plasma membrane.

To scrutinize whether sodium cations or chloride anions in NaCl were required to promote ferroptosis, we used N-methyl-d-glucamine chloride (NMDG-Cl) or sodium gluconate (Na-Glu), respectively, to replace NaCl in culturing media^32,38,39^. Intriguingly, depletion of [Cl^-^]_o_ in DMEM (Dulbecco’s Modified Eagle Medium) or Leibovitz’s L-15 media suppressed ferroptotic cell death upon RSL3 treatment, yet similar replacement of extracellular sodium cations cannot manifest comparable effect (Fig.2e-f). Flow cytometry analyses of cells stained with C11-BODIPY showed that a significant portion of RSL3-induced cells in [Cl^-^]_o_-depleted media possessed low level of lipid peroxidation (Fig.2g-h). Since the trypsin digestion and centrifugation processes before flow cytometry analyses may lower the percentage of fragile cells, we thus conducted microscopy analyses of *in situ* ferroptotic cells stained with C11-BODIPY. Remarkably, there was a significant higher portion of ferroptotic cells in [Cl^-^]_o_-depleted media accumulated high level of lipid peroxidation but did not undergo membrane rupture and cell lysis (Fig.2i). Lastly, we used a chloride ion reporter, (N-(Ethoxycarbonylmethyl)-6-methoxyquinolinium bromide (MQAE), which shows low fluorescence intensity in the presence of chloride ions, to monitor the Cl^-^ flow during ferroptosis. Consistent with a role of Cl^-^ in ferroptosis, we observed that upon RSL3 treatment, there was a decreased intracellular chloride ([Cl^-^]_i_) in cells and followed by an increased [Cl^-^]_i_ prior to plasma membrane rupture (Fig.2j). Taken together, our data showed that Cl^-^ is important for ferroptosis and there is Cl^-^ flow across the ferroptotic plasma membrane that happens concomitantly with water influx and membrane rupture.

### High [Cl^-^]_o_ is sufficient for eliciting ferroptosis

Having established that a low level of [Cl^-^]_o_ impedes ferroptosis of cells, we next studied the ferroptotic fate of cells in the presence of ectopically high level of [Cl^-^]_o_. Surprisingly, addition of high concentration of NaCl to the culturing media without any ferroptotic inducers elicited cell death that can be suppressed by canonical ferroptotic inhibitor Fer-1 (Fig.3a). The cell death induced by high NaCl in the media could be observed in multiple cell types and more importantly, was not suppressed by other programmed cell death inhibitors, suggesting that high NaCl does not cause a general cell death response but rather induces ferroptosis specifically (Fig.3b-c).

**Fig. 3.**
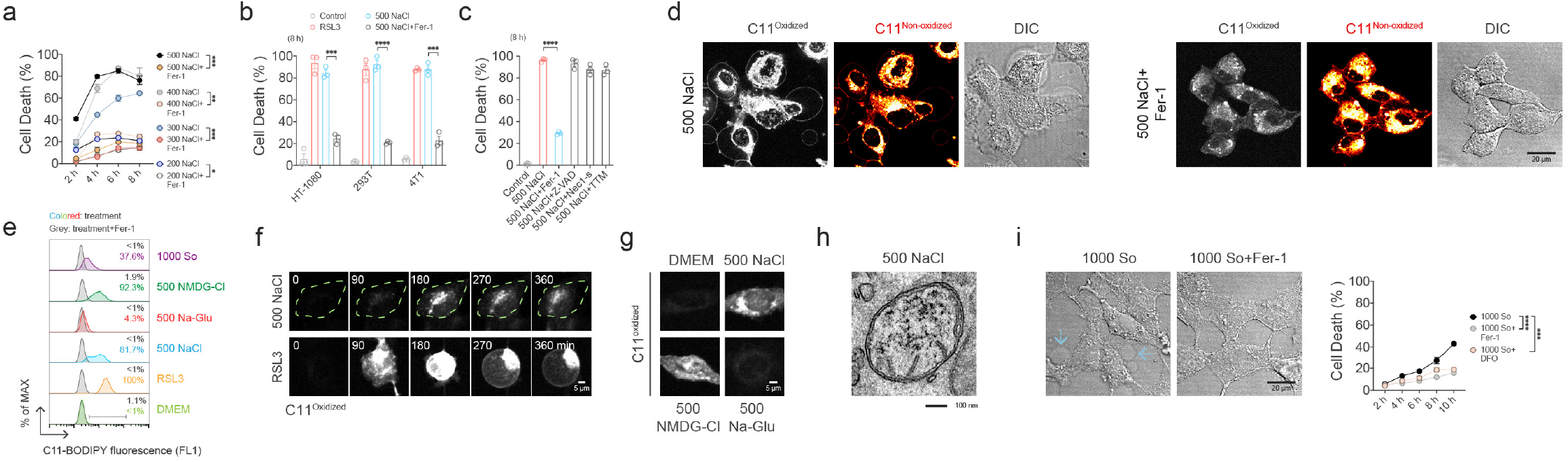
High [Cl^-^]_o_ is sufficient for eliciting ferroptosis. **a,** High NaCl elicits ferroptosis without other canonical ferroptotic inducers in a time and concentration dependent manner. **b,** High NaCl elicits ferroptosis in different cell types. **c,** High NaCl-induced cell death is suppressed by ferroptotic inhibitor Fer-1, but not by other types of programmed cell death inhibitors. **d,**500 mM NaCl increases lipid ROS level which can be inhibited by Fer-1. High contrast was adopted to visualize membrane swelling in the oxidized C11-BODIPY images. **e,** Lipid ROS production of different replacement conditions in DMEM media assessed by flow cytometry using C11-BODIPY. All treatments were assayed at 2 h. 500 NaCl represents DMEM media with extra 500 mM NaCl. 500 Na-Glu represents DMEM media with extra 500 mM Na-Glu. 500 NMDG-Cl represents DMEM media with extra 500 mM NMDG-Cl. 1000 So represents DMEM media with extra 1000 mM sorbitol. **f,** Lipid ROS production of 500 mM NaCl in DMEM media assessed by time-lapse confocal microscopy using C11-BODIPY. Dashed line indicates the cell boundaries. **g,** Lipid ROS production of different replacement conditions in DMEM media assessed by confocal microscopy using C11-BODIPY. **h,** Transmission electron microscopy image of a representative mitochondria from a 500 mM NaCl-treated cell. **i,**1000 mM sorbitol does not induce comparable percentage of ferroptotic cell death. Blue arrows indicate small membrane bubbles in the 1000 mM sorbitol treatment. The small bubbles are suppressed by Fer-1.

Similar to other ferroptotic inducers, high level of NaCl resulted in accumulation of lipid peroxidation that can be lowered by Fer-1 (Fig.3d-e). Interestingly, lipid peroxidation of the cell was specifically associated with the chloride ions in NaCl as cells grown in high Na-Glu media did not accumulate oxidized C11-BODIPY as compared to cells grown in high NMDG-Cl media (Fig.3e). Time-lapse microscopy analyses further confirmed that cells accumulated oxidized C11-BODIPY when cultured in high NaCl media containing chloride ions (Fig.3f-g). Additionally, electron microscopy analyses showed that cells grown in high NaCl manifested rounding mitochondria and disruption of mitochondria cristae, which are similar to typical features of ferroptosis (Fig.3h). Moreover, induction of ferroptosis by high NaCl was not simply due to the hyperosmotic stress on cells as high sorbitol with similar osmolarity could not induce comparable ferroptotic cell death rates (Fig.3i). Collectively, our data revealed a surprising effect of high [Cl^-^]_o_ in inducing ferroptosis.

### Cl^-^ acts downstream of lipid peroxidation

We have shown that Cl^-^ flow happens during ferroptosis and [Cl^-^]_o_ is playing a key role in driving ferroptosis. Since Cl^-^ contributes to electrochemical gradient across the plasma membrane and regulates the membrane potential, we speculated that there was a change in the membrane potential when cells undergo ferroptosis. To test if this was the case, we stained RSL3-treated cells with a membrane potential reporter bis-(1,3-dibutylbarbituric acid) trimethine oxonol (DiBAC4(3)) and observed that there was an increase of the fluorescence intensity as a function of time, indicating plasma membrane depolarization upon lipid peroxidation caused by GPX4 inhibition (Fig.4a). Consistently, addition of high NaCl to culture media resulted in plasma membrane depolarization, and [Cl^-^]_o_ was essential in this process (Fig.4b-c). We further found that plasma membrane depolarization by lipid peroxidation upon GPX4 inhibition was likewise affected by [Cl^-^]_o_ in the media (Fig.4d). Collectively, these data implicated that the membrane potential is involved in ferroptosis.

**Fig. 4.**
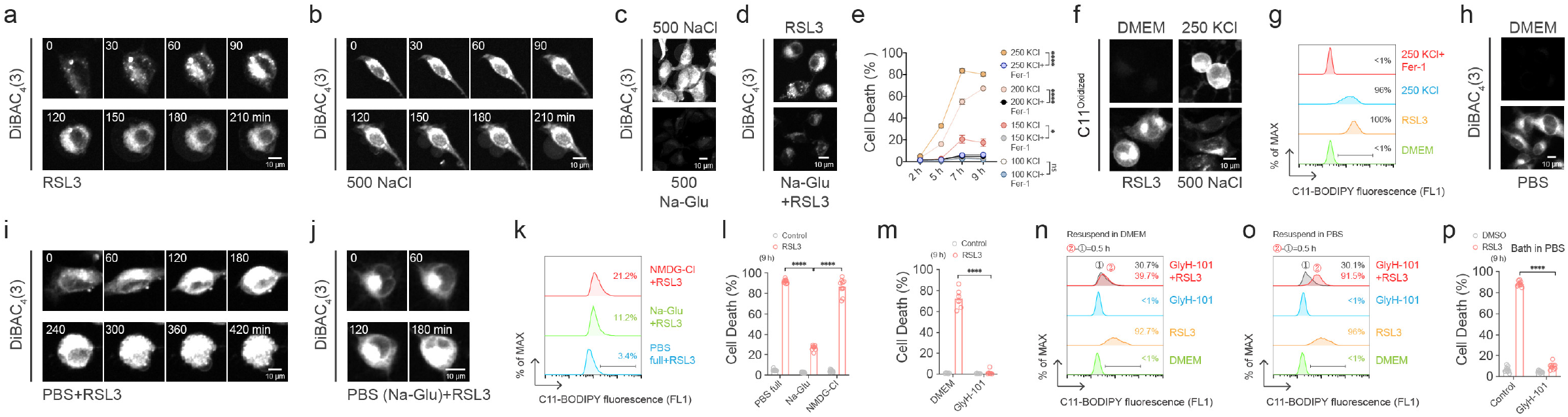
Cl^-^ acts downstream of lipid peroxidation. **a,** Membrane depolarization occurs in a RSL3-treated cell detected by DiBAC4(3). **b,** Membrane depolarization occurs in a 500 mM NaCl-treated cell detected by DiBAC4(3). **c,** Replacement of the chloride ion of 500 mM NaCl impairs depolarization. **d,** Replacement of the chloride ion of NaCl in DMEM impairs RSL3-induced depolarization. **e,** High KCl induces ferroptosis. KCl concentration, mM. **f,** Lipid ROS production of cells (250 mM KCl vs. 500 mM NaCl) assessed by confocal microscopy using C11-BODIPY. **g,** Lipid ROS production of 250 mM KCl-treated cells assessed by flow cytometry using C11-BODIPY. All treatments were assayed at 2 h. **h,** PBS induces membrane depolarization detected by DiBAC4(3). **i,** Membrane depolarization occurs in a RSL3-treated cell bathing in PBS detected by DiBAC4(3). **j,** Membrane depolarization occurs in a RSL3-treated cell bathing in Cl^-^-deprived Na-Glu-substituted PBS detected by DiBAC4(3). **k,** Lipid ROS production of ferroptotic cells in substituted PBS. 0.2 μM RSL3 was used in this experiment. All treatments were assayed at 1.5 h. **l,** Cl^-^-deprived PBS blocks ferroptosis. **m,** GlyH-101 blocks ferroptosis. **n,** Lipid ROS production of RSL3+GlyH-101 cells assessed by flow cytometry using C11-BODIPY. After 2 h of RSL3+GlyH-101 treatment in DMEM, cells were digested and resuspended in DMEM, in which lipid ROS level of RSL3+GlyH-101 cells did not change over time. **o,** Lipid ROS production of RSL3+GlyH-101 cells assessed by flow cytometry using C11-BODIPY. After 2 h of RSL3+GlyH-101 treatment in DMEM, cells were digested and resuspended in PBS, in which lipid ROS level of RSL3+GlyH-101 cells changed significantly over 0.5 h. **p,** Cell death analyses of RSL3+GlyH-101 cells bathed in PBS.

To test if changes of the membrane potential could influence ferroptosis, we cultured cells in media containing high KCl as the potassium ions are one of the keys in regulating the membrane potential (Fig.4e). We observed that higher number of cells underwent cell death when grown in an increased concentration of extracellular KCl. Cell death could be suppressed by Fer-1, indicating that changes of the membrane potential promote ferroptotic cell death. More importantly, KCl overload caused greater cell death compared to NaCl overload at similar concentrations, implying that potassium ions per se can potentiate ferroptosis (Fig.4e vs. Fig. 3a). Consistently, cells upon treatment of high KCl accumulated high level of oxidized C11-BODIPY (Fig.4f-g). Thus, membrane depolarization may favor the accumulation of lipid peroxidation.

Our aforementioned results showed that the [Cl^-^]_o_-depleted ferroptotic cells fall into two populations: cells with low lipid peroxidation; cells with high lipid peroxidation but not cell swelling or cell rupture. It is therefore hard to distinguish whether Cl^-^ plays upstream or downstream of lipid peroxidation. Since blockage of Cl^-^ flow may cause membrane hyperpolarization^40,41^, which may in turn limit the level of lipid peroxidation, we sought a method to force membrane depolarization on cells upon [Cl^-^]_o_ depletion. Interestingly, we found the PBS buffer depolarizes cells (Fig.4h). RSL3-induced ferroptotic cells possess higher DiBAC_4_(3) fluorescence in PBS buffer than cells in DMEM (Fig.4i vs. 4a). Upon NaCl substitution with Na-Glu, the substituted PBS retained its depolarization potency (Fig.4j). Intriguingly, [Cl^-^]_o_-depleted ferroptotic cells survive with a high level of lipid peroxidation (Fig.4k-l). Together, these data suggested that Cl^-^ acts downstream of lipid peroxidation.

Since the ion transport across the plasma membrane can be mediated by ion channels, we speculated that ion channels were involved in ferroptosis by regulating membrane potential. To this end, we found a chloride ion channel blocker, GlyH-101, can significantly inhibit ferroptosis (Fig.4m). When treated with RSL3, flow cytometry analyses showed that GlyH-101 treated ferroptotic cells bathing in DMEM possess relatively low level of oxidized C11-BODIPY compared to RSL3 treatment alone (Fig.4n). Since GlyH-101 treatment can hyperpolarize cells^40,41^, we again examined lipid peroxidation under the depolarizing PBS condition. When GlyH-101 treated ferroptotic cells were forcefully depolarized by PBS, we observed a significant increase of cells with lipid peroxidation over time in the presence of RSL3 (Fig.4o). Importantly, ferroptotic cell death remained significantly suppressed by GlyH-101 when bathed in depolarizing PBS, indicating that the chloride ion channel blocker GlyH-101 can abrogate ferroptosis while preserving a high level of lipid peroxidation (Fig. 4p).

## Discussion

Our study showed that the flow of Cl^-^ and water across the plasma membrane play an important role in regulating ferroptosis. Upon inhibition of GPX4, lipid peroxidation of the plasma membrane is accompanied by the membrane depolarization and increased Cl^-^ flow across the plasma membrane. Cl^-^ flow concomitantly drives water flow across the plasma membrane and results in cell swelling and membrane rupture, eventually leading to ferroptotic cell lysis.

Ferroptotic cells become quasispherical and swell as giant bubbles during the late phase before cell lysis. It is unclear how precisely ferroptosis culminates in cell swelling and lysis, despite enormous translational relevance. Previous ferroptosis studies have converged on machineries regulating lipid peroxidation, but how exactly lipid peroxidation triggers the eventual ferroptotic plasma membrane rupture is unknown^42^.

It remains enigmatic whether ferroptosis is mediated by a downstream pathway and/or a protein-controlled machinery. It was assumed that lipid peroxidation directly disrupts plasma membrane integrity leading to plasma membrane rupture during ferroptosis. However, our study provides evidence that cells could accumulate high level of lipid peroxidation but do not necessarily undergo cell lysis during ferroptosis. We find that the flow of Cl^-^ and water across the plasma membrane is necessary to drive plasma membrane rupture in the presence of lipid peroxidation. Typically, a steep spatial gradient of [Cl^-^]_o_ is maintained across the plasma membrane. When Cl^-^ flows into cells, the imbalance of ions across the plasma membrane causes a surge of osmolarity and expeditious cell swelling. Net gain of Cl^-^ and subsequent water influx are the driving forces for ferroptotic cell swelling.

Lipid peroxidation upon GPX4 inhibition is accompanied by the flow of Cl^-^ across the plasma membrane, which could be mediated by Cl^-^ channels. Whether Cl^-^ channels are activated directly or indirectly by the lipid ROS is currently unclear, but fenestration exists within protein channels and channels could be opened by PUFA oxidation^43,44^. Furthermore, the flow of Cl^-^ also influences the membrane potential, which changes during ferroptosis. Depolarization of the plasma membrane, in the presence of GPX4 inhibition, promotes lipid peroxidation. Phosphate-buffered salines, which strongly depolarize the cell membrane, is frequently used as a bathing buffer to measure the level of lipid peroxidation by C11-BODIPY dye. Thus, caution should be taken in future when measuring lipid ROS level in ferroptotic studies.

We demonstrate an important role of chloride ions in NaCl in modulating ferroptosis. Hyperosmotic stress is involved in diseases and NaCl is frequently utilized to generate hyperosmotic stress^45–47^. A recent study using high NaCl to introduce hyperosmotic stress on cells showed that ferroptosis is promoted under hypertonicity^48^. Since high level of [Cl^-^]_o_ promotes ferroptosis, the use of high NaCl to introduce hyperosmotic stress should be cautious. Our study used poorly permeable sorbitol or sucrose as solutes to introduce hyperosmotic stress, thus could better distinguish the contributions of NaCl and hyperosmolarity to ferroptosis.

Conceivably, the application of membrane potential inhibitors or activators as well as genetic manipulation of Cl^-^ flow could be a useful therapeutic method for treating ferroptosis related diseases. Our study offers a rethinking of ferroptosis execution mechanism and suggests a new paradigm that could be targeted for ferroptosis studies.

## Methods

### Cell culture

HT-1080 fibrosarcoma cells (obtained from the Shanghai Institute of Cell Biology, Chinese Academy of Sciences) were maintained and passaged every 1–3 days in Dulbecco’s modified Eagle’s medium (DMEM, Gibco) supplemented with 10% fetal bovine serum (FBS, ExCell Bio.) and 1% penicillin/streptomycin (Beyotime), in a humidified incubator at 37°C with 5% CO_2_. HEK293T and 4T1 cells were purchased and maintained as HT-1080 cells. NaCl-depleted media were custom made from Sunncell Bio. Then chloride or sodium was replaced with equal molarity sodium gluconate or NMDG-Cl. In some experiments where gradients of NaCl were indicated, the osmolarity was maintained by addition of sorbitol. NaCl overloaded media were made by adding the specified amount of NaCl powder into the basic cell culture media. Unless otherwise specified, all experiments were done in DMEM and HT-1080 cells and RSL3 final concentration is 2 μM.

### Chemicals and reagents

The following reagents were used at the indicated concentrations: Ferrostatin-1 (Fer-1, MCE, #HY-100579) 5-10 μM; Deferoxamine mesylate (DFO, MCE, #HY-B0988) 100 μM; Z-VAD (also named Z-VAD-FMK, Glpbio, #GC12861) 20 μM; Necrostatin 2 racemate (Nec1-s, Selleck, #S8641) 30 μM; Ammonium tetrathiomolybdate (TTM, Macklin, #15060-55-6); (1S,3R)-RSL3 (Selleck, #S8155) 0.5-2 μM; Erastin (Sigma, #E7781) 10 μM; C11-BODIPY(581/591) 5 μM (Invitrogen, #D3861). Sorbitol (#S1876); Sucrose (Macklin, #S818045); GlyH-101 (MCE, # HY-18336) 5 μM.

### Cell death assay

Cells were seeded in 96-well microplates at appropriate cell density and incubated overnight at 37°C in a humidified tissue culture incubator containing 5% CO_2_. Following 0-24 h treatments, cells were stained with hoechst 33342 (1 μg/mL, Beyotime, #C1022) and propidium iodide (PI, #ST511, 10 μg/mL) to monitor total cell number and dead cell number, respectively. Fluorescence was read by a cell imaging multimode fluorescence plate reader at indicated time points. Percentage of cell death was calculated as PI positive cell number over total Hoechst positive cell number. In some experiments, cell death was also verified by LDH assay (Beyotime, #C0017) following the manufacturer’s instructions.

### Lipid ROS assay using flow cytometry

Cells were seeded in 12-well microplates at appropriate cell density and were incubated overnight at 37°C in a humidified tissue culture incubator containing 5% CO2. The cell culture medium was changed with medium (DMEM or L-15) containing 5 μM C11-BODIPY dye and the corresponding drugs (RSL3 and or inhibitors). Microplates were placed back in the tissue culture incubators. After culturing for the indicated time, media were discarded and cells were harvested by trypsinization. Centrifugation was applied and cells were resuspended in DMEM or PBS containing 5 μM C-11 BODIPY dye and the corresponding drugs (RSL3 and or inhibitors). Lipid ROS level was assessed using the CytoFLEX Flow Cytometer (Beckman) with a 488 nm laser on an FL1 detector. Data were analyzed using FlowJo. In each sample, ~10,000 cells were analyzed.

### Microscopy

Cells were seeded onto 4-chamber 35 mm glass-bottomed dishes (tissue culture-treated, JingAn Bio, China) at appropriate cell density overnight. After changing media with drugs, cells were maintained in a 37°C chamber with 5% CO_2_ during the entire imaging process. Confocal microscopy systems were utilized for capturing fluorescent images. Each chamber was imaged at 2 locations during each time point. Time interval is 15 minutes. For C11-BODIPY live cell imaging, low laser power should be used to avoid phototoxicity and photoconversion. Band pass (recorded emission wavelength: 485-565 nm) was used for collecting oxidized C11-BODIPY signals. In all movies, 0 min indicates the start of movie. There could be a 20-60 min gap between changing media and the start of movie.

### MQAE

Cells were seeded onto 4-chamber 35 mm glass-bottomed dishes (tissue culture-treated, JingAn Bio, China) at appropriate cell density overnight. 5 mM MQAE was added into each chamber and 405 nm laser was used for time-lapse imaging.

### DiBAC_4_(3)

Cells were seeded onto 4-chamber 35 mm glass-bottomed dishes (tissue culture-treated, JingAn Bio, China) at appropriate cell density overnight. 1 μM DiBAC4(3) was added into each chamber and FITC filter set (recorded emission wavelength: 450-700 nm) was used for time-lapse imaging.

### Transmission electron microscopy

The back of a the sapphire substrate was marked to distinguish the top or the down sides. The sapphire substrate was soaked in 5% polylysine solution for 5 min and was dried in a 50°C incubator for 30 min. The sapphire substrate was placed into a six-well plate. HT-1080 cells were seeded onto the sapphire substrate with appropriate cell density and cultured in a 37°C, 5% CO_2_ cell culture incubator for 24 h. After 500 mM NaCl treatment, cells and the sapphire substrate were fixed together in paraformaldehyde for 2 h at room temperature, and subsequently at 4°C overnight. Cells were fixed with 3% glutaraldehyde (SPI-CHEM Inc., USA) at 4°C for 4 h, washed with sodium cacodylate buffer 3 times at an interval of 2 h. Then cells were soaked in 1% osmic acid (SPI-CHEM Inc., USA) for 2 h at 4°C and washed with 0.1 M sodium cacodylate buffer twice with 15 min washing each time. Cells were stained with saturated uranyl acetate dye (SPI-CHEM Inc., USA) for 2 h at room temperature. For dehydration and infiltration, cells were incubated in 50% alcohol for 10 min at 4°C, 70% alcohol for 10 min at 4°C, 80% alcohol for 10 min at room temperature, 90% alcohol for 15 min at room temperature, 100% alcohol twice for 10 min at room temperature. Cells were infiltrated twice with acetone for 15 min at room temperature. Then cells were infiltrated with complete embedding solution and acetone (1:1) for 3 h at room temperature. The complete embedding solution was Eponate12 epoxy resin (Ted Pella, Inc., USA). Cells were Infiltrated with complete embedding solution and acetone (1:2) for 3 h at room temperature and soaked in complete embedding solution overnight at room temperature. Then cells in complete embedding solution were transferred to the incubator at 40°C for 12 h and subsequently 60°C for 48 h. UC7 ultramicrotome (Leica Microsystems Ltd., Germany) was used for ultrathin section with section thickness of 90 nm. JEM-1400 transmission electron microscope (JEOL Ltd., Japan) was used for image capturing with operating voltage of 80 kV. Images were acquired with 832 digital micrograph software (Gatan, Inc., USA).

### Quantification and statistical analysis

Statistical analysis was performed using an unpaired Student’s t-test. All summarized data are reported as means ± SEM (*P < 0.05, **P < 0.01, ***P < 0.001, ****P < 0.0001. ns, not significant). Analyses were performed using GraphPad Prism 6 software. Linear regression lines were performed using inbuilt algorithms (simple linear regression).

## Acknowledgements

We would like to express our most sincere gratitude to all mentors from our group (Leader: Dr. Yu Lan) of the Program for Guangdong Introducing Innovative and Entrepreneurial Teams. We would like to sincerely thank Dr. Jiekai Chen, Dr. Ronghan He and Mr. Jiewen Kong for supporting the project, Ms. Ziqi Wang for her supports and specialties in helping with the schematics, Dr. Huabin Wang for proofreading the manuscript. We sincerely thank our families and friends for their supports in difficult times. We would like to dedicate this work with eternal gratitude to the memory of Fan Zhou for his invaluable help and love in doing Science. Funding derived from the Program for Guangdong Introducing Innovative and Entrepreneurial Teams, grant number 2017ZT07S347; the Guangzhou Basic and Applied Basic Research Foundation, grant number 202102020509; the National Natural Science Foundation of China, grant number 32270770; the National Natural Science Foundation of China, grant number 31701174.

## Author Contributions

JH conceived and initiated the project. JH, TGC and XZ designed and supervised the project. JH, TGC and XZ discussed and wrote the manuscript. DH, LC initiated the chloride direction. ZD, DH initiated the membrane potential direction. CW, DH, ZD carried out the genetic manipulation experiments. RH, JZ, SZ, TB, YC contributed novel ferroptosis inhibitors. HM, KX helped with the microscopy imaging. SW and ZD conducted the electron microscopy experiments.

## Data availability

All data underpinning the conclusions in this work are available from the corresponding authors upon reasonable request.

